# Benefit-cost analysis of the policy of mandatory annual rabies vaccination of domestic dogs in rabies-free Japan

**DOI:** 10.1101/448258

**Authors:** Nigel C. L. Kwan, Akio Yamada, Katsuaki Sugiura

## Abstract

Japan is one of the few rabies-free countries/territories which implement the policy of mandatory vaccination of domestic dogs. In order to assess the economic efficiency of such policy in reducing the economic burden of a future canine rabies outbreak in Japan, a benefit-cost analysis (BCA) was performed using probabilistic decision tree modelling. Input data derived from simulation results of published mathematical model, field investigation conducted by the authors at prefectural governments, literature review, international or Japanese database and empirical data of rabies outbreaks in other countries/territories. The current study revealed that the annual costs of implementing the current vaccination policy would be US$160,472,075 (90% prediction interval [PI]: $149,268,935 – 171,669,974). The economic burden of a potential canine rabies outbreak in Japan were estimated to be US$1,682,707 (90% PI: $1,180,289 – 2,249,283) under the current vaccination policy, while it would be US$5,019,093 (90% PI: $3,986,882 – 6,133,687) under hypothetical abolition of vaccination policy, which is 3-fold higher. Under a damage-avoided approach, the annual benefits of implementing the current vaccination policy in expected value were estimated to be US$85.75 (90% PI: $55.73 – 116.89). The benefit-cost ratio *(BCR)* was estimated to be 5.35 × 10^−7^ (90% PI: 3.46 × 10^−7^ – 7.37 × 10^−7^), indicating that the implementation of the current policy is very economically inefficient for the purpose of reducing the economic burden of a potential canine rabies outbreak. In worse-case scenario analysis, the *BCR* would become above 1 (indicating economic efficiency) if the risk of rabies introduction increased to 0.04 corresponding to a level of risk where rabies would enter Japan in 26 years while the economic burden of a rabies outbreak under the abolition of vaccination policy increased to $7.53 billion. Best-case analysis further revealed that the economic efficiency of the current policy could be improved by decreasing the vaccination price charged to dog owners, relaxing the frequency of vaccination to every two to three years and implementing the policy on a smaller scale, e.g. only in targeted prefectures instead of the whole Japan.

## 1. Introduction

Since 1958, Japan has been free from animal rabies, i.e. classical rabies virus (RABV), and is one of the few rabies-free countries or territories (under the standards of World Organisation for Animal Health [OIE]) which still implement the policy of mandatory vaccination of domestic dogs [1]. Under the Rabies Prevention Law enacted since 1950, the policy of registration and vaccination of domestic dogs against rabies is enforced by the health departments of prefectural governments under the order of Ministry of Health, Labour and Welfare (MHLW) [1]. Pet owners in Japan are obliged to vaccinate their dogs against rabies every year either by attending private animal clinic anytime during the year or a vaccination campaign organised by prefectural governments and Japan Veterinary Medical Association (JVMA) during April to June. Each year the respective prefectural government would assign the duty and provide a fund to the local Veterinary Medical Association to organise the rabies vaccination campaign mentioned above in multiple cities within the prefecture. In the decade 2007–2016, the reported national vaccination rate averages 73.1%, but the actual vaccination coverage is estimated to be 43.2% when adjusted for an average estimated registration rate of 59.2% during the same period [2, 3].

The current risks of rabies re-introduction into Japan have recently been assessed as very low and it would take on average 49,444 years until the introduction of one rabies case through the importation of dogs and cats worldwide due to a strict import regime managed by the Ministry of Agriculture, Forestry and Fisheries of Japan (MAFF) [4, 5]. Further, mathematical simulation model predicted a very low risk of local spread, if rabies were to be introduced into Japan, as the mean outbreak size was estimated to be 3.1 and 4.7 dogs in Hokkaido and Ibaraki Prefectures, respectively [6]. Together with the low owner compliance mentioned above, massive debate has been raised in the country over whether the current annual rabies vaccination policy in domestic dogs should be maintained.

The main advantage of implementing a pre-emptive vaccination policy in a rabies-free setting is that it facilitates a pre-existing herd immunity which could lessen the magnitude or impact of an introduced outbreak. For canine rabies-endemic countries/territories, the World Health Organization (WHO) recommends a minimum 70% vaccination coverage in domestic dog population as the most cost-effective control measure [7]. In contrast, there is no international standard or well-established guideline on the prevailing conditions that should prompt a rabies-free country to implement a pre-emptive vaccination policy [8]. Indeed, major rabies-free countries including United Kingdom, France, Australia and New Zealand are not relying on domestic dog vaccination as a preventive strategy because it is perceived that early detection of suspected rabid animals and an immediate response to contain the outbreak are most important. On the other hand, Hong Kong, a rabies-free territory since 1988, implements the policy of mandatory dog vaccination every three years due to substantial risk of rabies being introduced from China. Malaysia also adopts a dog vaccination policy building a 50 to 80 km immune belt along the border with Thailand; the country had been declared rabies-free since 2012 until 2015 when the disease was introduced most probably from Thailand [9]. Taiwan enforces rabies vaccination in both domestic dogs and cats, and had been rabies-free since 1961 until an epizootic of Chinese ferret-badger started in 2013 [10].

Benefit-cost analysis (BCA) is an important tool at both a global and national level that illustrates the benefits of disease management projects per dollar spent and determines the economic efficiency of alternative management actions [11]. BCA on the control interventions against various animal diseases have been conducted, particularly on the oral vaccination against wildlife rabies [12, 13]. The current study aimed to perform a BCA using decision tree modelling to assess the economic efficiency of the current annual rabies vaccination policy in domestic dogs in Japan and serve as a pilot study providing scientific insight into the rationale behind the maintenance of such policy.

## Materials and Methods

### Decision tree model and cost estimation framework

A stochastic decision tree model was constructed comparing two strategies: 1. under the current annual vaccination policy a rabid dog is introduced into Japan resulting in an outbreak and 2. under hypothetical abolition of vaccination policy a rabid dog is introduced into Japan resulting in an outbreak with greater impact (Fig 1). The annual number of dogs and cats imported into Japan worldwide (majority through airplane and a few through ship) is approximately 8,000 and 2,000 through Animal Quarantine Service (AQS) managed by MAFF and United States Force Japan (USFJ), respectively. Therefore, the annual probability of rabies introduction into Japan through these pathways reported previously *(P_annual),* i.e. 2.57 × 10^−5^, was input into the relevant chance nodes of the decision tree, while the effect of any increase in the risk of rabies introduction, e.g. illegal importation, was tested in scenario analysis described below [5]. There are potentially other rabies entry pathways, e.g. via fishing boat, passenger ferry and shipping containers. However, it was assumed that the risk of introduction would not increase significantly even if the base model took into account the risk of introduction through these pathways, which is a reasonable assumption considering the very small number of dogs and cats imported through these pathways and the results of a previous quantitative risk assessment [4]. Finally, it should be noted that rabies introduction via the land route was not considered given that Japan is geographically isolated by the sea. The time horizon of the model was one year and no discount rate was applied to the calculations of benefits and costs. The monetary values reported in the current study were based on the exchange rate of 1 US dollar = 112.17 Japanese yen (2017 World Bank data).

**Fig.1.**
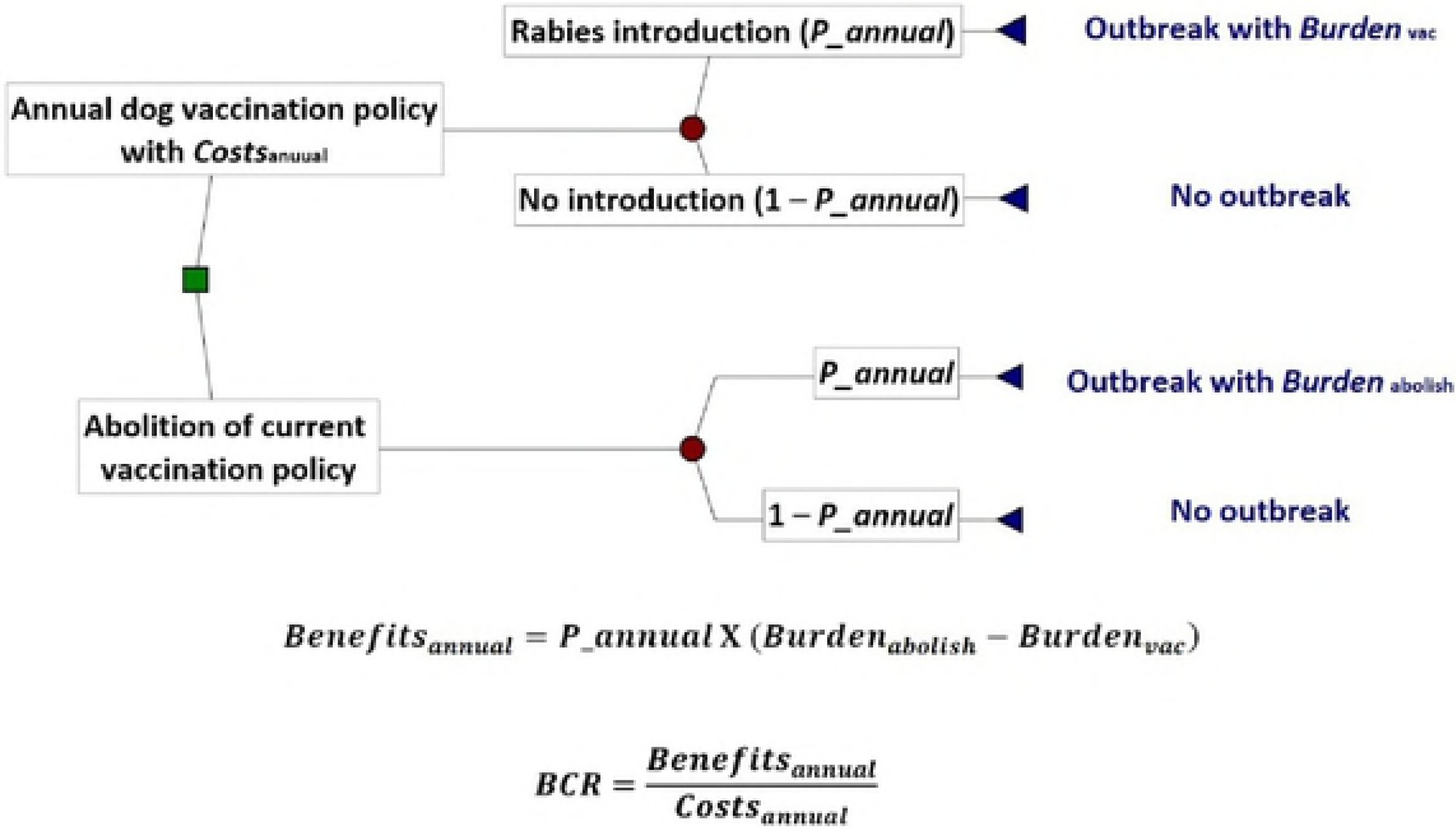
Conceptual framework of the current benefit-cost analysis through decision tree modelling – the economic efficiency of implementing the current dog vaccination policy for the purpose of reducing the economic burden of a potential canine rabies outbreak in rabies-free Japan is indicated by the benefit-cost ratio, *BCR.*

The present BCA adopted a societal perspective where the benefits and costs of maintaining the current annual rabies vaccination policy were considered for every relevant stakeholder in the community. In essence, such policy was considered to benefit everyone in the community since it could reduce the impact of the outbreak (in terms of both the number of cases and the duration), hence decreasing the economic burden of the outbreak in terms of lower costs in implementing rabies control measures for the government and lower risk of contracting rabies for the local people (which would lead to fewer people receiving medical treatment and a lower probability of human death). Hence, the benefits of maintaining the current rabies vaccination policy were calculated as incremental benefits using a damage-avoided approach described below. On the contrary, the costs of maintaining the current policy were considered to be borne by dog owners who vaccinate their dogs against rabies (note that they also bear the gross profits made by private animal clinic or JVMA) and the government in providing funds to JVMA to organize the vaccination campaign.

The annual costs of implementing the rabies vaccination policy and the economic burden of a canine rabies outbreak in Japan were estimated based on published frameworks with specific modifications to accommodate the rabies-free setting of Japan [14, 15]. The authors conducted three investigation trips to Ibaraki, Tokushima and Miyazaki Prefectures during 20 July 2016 to 29 September 2016 and interviewed the prefectural government officials to obtain necessary information regarding local rabies prevention system. Other data derived from extensive literature review, international or Japanese database and empirical data of rabies outbreaks in Asian countries/territories including Taiwan and Malaysia and European countries such as France and the Netherlands.

### Estimation of the annual costs of implementing the dog rabies vaccination policy in Japan (*Costs*_annual_)

#### Direct vaccination costs

The key characteristics of companion dog ownership in Japan are summarized in Table 2. In 2015, 4,688,240 companion dogs were vaccinated against rabies based on the official figures published by MHLW [2] and the number of owners involved was estimated to be 3,780,839 assuming one representative from each household with dog ownership. Approximately 47.2% to 48.6% of dog owners would attend the vaccination campaign based on information from the investigated prefectural governments. The price of single vaccination charged in vaccination campaign in the 47 prefectures of Japan ranges from $20.50 to $27.64. The average single vaccination price was estimated to be $24.61 based on a weighted mean of all the vaccination prices accounting for the number of registered dogs vaccinated in each prefecture. In addition, owners would need to pay $4.90 for a tag certifying the dog’s ! vaccination. Overall, the direct cost of a single dog rabies vaccination (*Cost*_vac_) was set as $29.52 assuming this proxy includes vaccine cost, material costs such as i needle, syringe and alcohol swab, overhead costs (including staff salaries and administrative cost), logistic costs and gross profit. It should be noted that the value of *Cost*_vac_ excluding gross profits would correspond to the unit value of the funds *’* provided by the government to JVMA for organization of annual vaccination campaign, i.e. the total amount of funds divided by the number of dogs vaccinated in campaign each year. The direct cost of single dog rabies vaccination at animal clinic) was also set as *Cost*_vac_ based on the observation that the dog rabies vaccination price charged in annual campaign would be very similar to average price charged in animal ! clinic in each respective prefecture, e.g. at Tokyo Metropolis the vaccination price (excluding the price of certification tag) was $27.64 in campaign, while the median price was $26.25 in animal clinics (n=1,365) based on 2015 Tokyo Veterinary Medication Association survey data.

#### Indirect costs

Indirect costs included opportunity costs of time and transport costs for owners and advertisement costs for the government. The opportunity costs of time were estimated using the human capital approach based on productivity or income loss. Such loss was calculated by weighing the number of working days lost by the daily gross domestic product (GDP) per capita. No transport cost was considered for owners attending vaccination campaign assuming that the majority of them would either walk or ride the bicycle based on information from the investigated prefectural governments. For owners who attend an animal clinic for vaccination, it was assumed half of them would drive and the other half would walk. Advertisement costs were considered for the production of publicity materials such as posters and leaflets by MHLW.

### Estimation of the economic burden of a canine rabies outbreak in Japan

The current model predicted the economic burden of a hypothetical canine rabies outbreak in Ibaraki Prefecture under the current vaccination policy with a coverage of 51.8% (*Burden*_vac_) and under the abolition of such policy, i.e. vaccination coverage of 0% (*Burden*_abolish_), respectively, according to the simulation results in Kadowaki et al. [6]. Ibaraki Prefecture was selected for investigation because it is representative of the current situation of dog ownership in Japan in terms of proportion of households with dog ownership, dog registration and rabies vaccination rates, companion dog density and dog-to-human ratio (Table 1). The main epidemiological characteristics of the simulated outbreak were considered in terms of the number of rabid dogs, i.e. mean outbreak size, and the duration of the outbreak, i.e. mean epidemic period – an outbreak would last for 68.2 days involving 4.7 rabid dogs under the current vaccination policy, while it would last for 152.5 days involving 21.7 dogs under the abolition of vaccination policy. It was assumed that the introduced rabies disease would not become endemic in the country and would come to an end under control interventions as predicted by the simulation model and according to the experiences in Western Europe [16]. Thus, the economic burden was considered on the basis of incurred expenses of a single rabies outbreak.

**Table 1.** Characteristics of companion dog ownership in Japan and Ibaraki Prefecture based on official figures in 2015.

| **<u>Characteristics</u>** | **<u>Japan</u>** | **<u>Ibaraki Prefecture</u>** |
| --- | --- | --- |
| **Proportion of households with dog ownership** | 14.42% | 17.52% |
| **Number of households with dog ownership<sup>1</sup>** | 7,985,000 | 196,986 |
| **Estimated number of companion dogs** | 9,917,000 | 244,263 |
| **Habitable area (km<sup>2</sup>)** | 122,631 | 3975 |
| **Companion dog density (dogs / km<sup>2</sup>)** | 81 | 61 |
| **Human population** | 127,094,745 | 2,917,000 |
| **Dog-to-human (thousands) ratio** | 78 | 84 |
| **Number of registered dogs reported by MHLW** | 6,526,897 | 176,628 |
| **Estimated dog registration rate** | 65.82% | 72.31% |
| **Number of dogs vaccinated against rabies reported by MHLW** | 4,688,240 | 118,387 |
| **Estimated rabies vaccination rate** | 47.27% | 48.47% |

<sup>1</sup> Average number of dogs owned by each household with dog ownership was 1.24 ( $n=50,000$ ) [3].

### Dog rabies control costs

Based on the national rabies contingency plan, it was assumed that the prefectural government rabies control team would respond to the outbreak by taking actions including epidemiological investigation (this involves contact tracing of all the dogs and other susceptible animals in close contact with the index case and identification of potential rabid dog-bite victims), emergency vaccination of dogs and depopulation (culling) of stray dogs around the outbreak area [17]. The cost of stray dog depopulation was calculated as an incremental cost since the capture and management of stray dogs and dogs without a registration tag and vaccination are already being conducted as part of the daily rabies prevention system.

### Human rabies prevention costs, i.e. post-exposure and pre-exposure prophylaxis (PEP and PrEP) costs

#### PEP due to rabid dog exposure

The number of human victims bitten or injured by a rabid dog was based on Kadowaki et al. [6]. It was assumed that all victims suffer either a Category II or III exposure requiring PEP under WHO recommendations and all of them can receive timely complete PEP given the fact that Japan is a developed country with a very high human development index of 0.903 [15, 18, Human Development Report 2015 data]. Currently there are two types of human rabies vaccine available in Japan, i.e. Japanese PCEC-K vaccine and imported vaccine such as Verorab and Rabipur [19]. An intramuscular 5-dose Essen regimen (which is the most common method used in local hospitals or clinics) with a fixed cost of $129 was considered for the direct medical cost of PEP, while the indirect costs were considered in a similar manner mentioned above [20, 2018 Tokyo Metropolitan Cancer and Infectious Disease Center Komagome Hospital data].

In terms of rabies immunoglobulin (RIG), it was assumed that human RIG and/or equine RIG would be imported for emergency use in case of a rabies outbreak (no such product is currently available in Japan) [19]. The number of patients with Category III exposure requiring RIG on Day 0 was estimated based on the data of outbreaks in the Netherlands and Greece: 50% (n=42) to 72% (n=96) of the patients receiving PEP would have a Category III exposure when bitten by a rabid dog [21, 22]. The efficacy of timely complete PEP was assumed to be 100% and so no human death, i.e. Years of Life Lost (YLL), was considered.

#### PEP due to public panic

In the early stage of the 2013 Taiwan epizootic, 5,335 persons injured with animal bites or scratches (78% due to a dog or cat) applied for free government-funded PEP during a 72-day period when 157 rabies cases were confirmed [23]. However, 35.5% of these applicants came from areas where no rabies case was reported, and only seven applications were ultimately proved to be caused by a rabid animal, i.e. Chinese ferret-badger. Since Japan has been rabies-free for over half a century, it can be foreseen that a rabies outbreak would cause massive public panic in the country leading to unnecessary use of PEP similar to the situation in the Taiwan epizootic. According to the Ministry of the Environment [24], the average daily incidence of dog-bite victims was reported to be 12 persons in 2016 (note that the number of victims exposed to animals other than dogs was not included in the calculation due to a lack of official data; for reference nine cases of fatal injury due to a cat was reported in 2016). Based on this, it was assumed that on a daily basis six persons (50% of the daily incidence) and ten persons (80% of the daily incidence) would receive PEP due to public panic during a rabies outbreak under the current vaccination policy and abolition of such policy, respectively. The direct and indirect costs involved were then estimated in a similar manner as described above, while the use of RIG was not considered for Category III exposure in this case.

#### Occupation PrEP

The original members of the prefectural government rabies control team and the diagnostic laboratories were assumed to have all received PrEP. Therefore, the direct costs of PrEP, using a three-dose intramuscular WHO regimen, were considered for the additional government officers who join the rabies control team in face of an outbreak [18, 25]. The indirect costs were assumed to be reflected in the labour costs of the officers and therefore were not considered.

#### Surveillance costs

Surveillance costs involve: 1. diagnostic testing of all rabid dogs, i.e. the positive cases, and 2. ongoing testing of all suspected animals during the outbreak and after the outbreak for two years to declare and verify rabies-free status according to OIE standards. In terms of the testing of suspected animals, the surveillance data of France was used as a proxy since the country has an intensified surveillance system due to regular rabies introductions, i.e. during 2008–2017, an average of five suspected animals were tested for rabies each day [16, 2017 Rabies Bulletin Europe data]. It was assumed that the level of active surveillance during a rabies outbreak in Japan under the abolition of vaccination policy would be the same as that in France, while it would not be as intensified during an outbreak under the current vaccination policy, i.e. five and three suspected animals are tested each day, respectively.

### Model implementation and outputs

The decision tree model (Fig 1) was developed in PrecisionTree and @Risk Version 7.5.1 (Palisade Corporation) within Microsoft^®^ Excel 2016, and was run with 5,000 iterations using Latin Hybercube sampling for each simulation. Information on cost data and input variables into the model is summarised in S1 Table.

Outputs of the model included the economic burden of a rabies outbreak in Japan under mandatory vaccination policy (*Burden*_vac_) and under abolition of vaccination policy (*Burden*_abolish_), respectively, and annual costs of implementing the current vaccination policy (*Costs*_annual_) were estimated. Utilising a damage-avoided approach, the annual benefits of implementing the current vaccination policy in expected value (*Benefits*_annual_) was calculated:

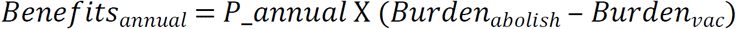

The benefit-cost ratio *(BCR)* was then given by: 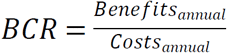

If the *BCR* is greater than 1, the implementation of the current annual vaccination policy is an economically-efficient strategy and vice versa (Shwiff et al., 2016).

### Sensitivity and scenario analyses

To assess the uncertainty in the current model, sensitivity analysis was conducted using Spearman’s correlation coefficient to rank all the input parameters according to their contributions to the variance in *BCR.*

The following three scenario analyses were performed to assess their effects on BCR:

1. Reduction in the direct medical cost of single dog rabies vaccination, i.e. *Cost*_vac_ – under the current policy, owners bear most of the costs including direct and indirect costs involved in bringing their dogs to annual rabies vaccination. The potential to reduce *Cost*_vac_ has been highlighted in certain prefectures where a few veterinary clinics charge a vaccination price as low as $8.92.
2. Worst-case scenario – this aimed to analyse the following two possible events: 1) the economic burden of a rabies outbreak under the abolition of vaccination policy, i.e. *Burden*_abolish_, was underestimated (relative to that under the current vaccination policy, i.e. *Burden*_vac_), 2) and the risk of rabies introduction into Japan increases in unforeseen circumstances, e.g. smuggling of animals. The parameters *Burden*_*abolish*_ and *P_annual* were increased in a stepwise fashion to model such situation. It should be noted that, by increasing the value of *Burden*_abolish_ (relative to *Burden*_vac_), this worse-case analysis would also indirectly address the effect of additional economic burden due to outbreak situations not considered in the current model, e.g. a rabies outbreak involving wildlife species or domestic cat;
3. Best-case scenario – this aimed to explore under what specific circumstances the economic efficiency of maintaining the current dog rabies vaccination policy could be maximized. *Costs*_annual_, by considering the number of companion dogs vaccinated with rabies, the frequency of vaccination and *Cost*_vac_, were decreased, while *Burden*_*abolish*_ and *P_annual* were increased to model the best-case situation. In particular, it has been highlighted that the current annual vaccination policy could be amended to one requiring less frequent boosters with the use of the domestic RC-HL strain vaccine, i.e. a booster is required within one year after primary vaccination and then every two to three years [26].

## Results

### Model outputs

Information on the simulated model outputs is summarised in Table 2 and Fig 2. The annual costs of implementing the current dog rabies vaccination policy were estimated to be $160,472,075 (90% prediction interval [PI]: $149,268,935 – 171,669,974). The economic burden of a rabies outbreak in Japan was estimated to be $1,682,707 (90% PI: $1,180,289 – 2,249,283) under the current vaccination policy, while it would be $5,019,093 (90% PI: $3,986,882 – 6,133,687) under the abolition of vaccination policy, which is 3-fold higher. The annual benefits of maintaining the current vaccination policy in expected value were estimated to be $85.75 (90% PI: $55.73 – 116.89). The benefit-cost ratio was estimated to be 5.35 × 10^−7^ (90% PI: 3.46 × 10^−7^ – 7.37 × 10^−7^).

### Sensitivity and scenario analyses

Result of sensitivity analysis is illustrated in Fig 3. The top five most uncertain parameters are the number of patients receiving PEP due to public panic under the abolition of vaccination policy (*N_panic,abolish_*), the daily number of suspected rabid animals tested under active surveillance under the abolition of vaccination policy (*N_d-survey,abolish_*) and under the current vaccination policy (*N_d-survey,vac_*), respectively, the number of patients receiving PEP due to public panic under the current vaccination policy (*N_panic,vac_* and the number of working days lost per owner per dog vaccination Results of scenario analysis are shown in Table 3 and 4 and Fig 4 and 5. The analysis of reduced direct medical cost of single dog rabies vaccination revealed that the *BCR* was 3.59 × 10^−6^ when *Cost*_vac_ was reduced to zero, highlighting that the implementation of the current annual vaccination policy would still still be economically inefficient if one only considered the indirect costs of vaccination for the dog owners (Fig 4). The worst-case scenario analysis demonstrated that the *BCR* (including the 90% PI) would become above 1 when the annual risk of rabies introduction into Japan *(P_annual)* and the economic burden of a rabies outbreak under the abolition of vaccination policy (*Burden*_abolish_) simultaneously increased 1500-fold (Table 3). The best-case scenario analysis further revealed that under the circumstance of a 100-fold increase in both *P_annual* and *Burden*_*abolish*_, the maintenance of a pre-emptive dog vaccination policy in rabies-free Japan could become economically efficient, i.e. mean *BCR* = 2.61, if it were implemented on a smaller scale, i.e. in only one of the 47 prefectures in Japan using Ibaraki Prefecture as an example, with a 3-fold decrease in *Cost*_vac_ to $8.92 at a frequency of every two to three years (Table 4).

## Discussion

The current study assessed the merit of implementing mandatory annual rabies vaccination in domestic dogs in Japan using benefit-cost analysis. The estimated values of benefit-cost ratio *(BCR)* were very low, i.e. well below 1, indicating that the implementation of the current vaccination policy in rabies-free Japan is very economically inefficient for the purpose of reducing the economic burden of a potential canine rabies outbreak. The annual costs of implementing such vaccination policy (*Costs*_annual_) were estimated according to the data on registered dogs reported by MHLW and might have been under-estimated since the average national registration rate during 2007–2016 is estimated to be only 59.2% [2, 3]. Nonetheless, it is anticipated that companion dogs which are not registered by their owners are less likely to be vaccinated regularly against rabies. The study by Hidano et al. [27] highlighted that companion dogs taken infrequently for walks are significantly less likely to be vaccinated against rabies in Japan. In addition, adverse drug reaction and vaccine wastage were expected to contribute to only a minor component of *Costs*_annual_ and hence were not considered [28].

The economic burden of a rabies outbreak in Japan, was estimated to be $1.69 million and $5.02 million, under the current vaccination policy and the abolition of such policy, respectively. Such level of burdens, although not directly comparable due to the differences in model framework, appear similar to the annual costs of rabies control in Flores Island, Indonesia which were estimated to be $1.12 million [29]. Further, the government of Taiwan have initially spent over $4.5 million for the support of contingency actions and procurement of human and animal rabies vaccines for the epizootic started in 2013 [10].

It should also be noted that the results of the current study could be generalized to other potential rabies situations in Japan, e.g. outbreaks primarily involving domestic cat, particularly stray cats which are twice as common as stray dogs [24], or wildlife species such as common racoon and red fox which are common in the country. If a non-canine rabies outbreak occurred in Japan, dog rabies control measures considered in the current model including emergency dog vaccination and stray dog depopulation and epidemiological investigation would still take place as part of the contingency plan, while the costs of PEP due to public panic would still be expected to constitute a considerable part of the economic burden as demonstrated in the Taiwan epizootic of Chinese ferret-badger [10]. On top of these basic components of the economic burden, there would be additional costs incurred in containing a non-canine rabies outbreak, e.g. extra manpower would be needed to support immediate stray cat depopulation in face of an outbreak involving domestic cat, while oral rabies vaccination (ORV) might be implemented in the long term if a wildlife rabies outbreak became endemic in the country [30]. Overall, the current study provided a generic framework for future research to estimate the potential economic burden of different rabies incursions in Japan.

To accommodate the unique situation in Japan, the current study did not consider certain components of the economic burden of canine rabies as suggested in Hampson et al. [15] and Knobel at al. [14]. Livestock losses were not included as significant losses were considered unlikely due to a single dog rabies incursion as indicated in historical incidence [31, 32]. In addition, the costs of potential human death, i.e. Years of Life Lost (YLL), were not considered as explained above, but it is possible that some patients with Category III exposure from a rabid dog could not receive timely rabies immunoglobulin (RIG) since it is currently not available in Japan. The reported probability of contracting rabies after Category III exposure to a rabid dog ranges between 0.03 and 0.25 when an incomplete post-exposure prophylaxis (PEP) without RIG is used [33]. Zhang et al. [34] have emphasized the importance of RIG in a PEP regimen and the inefficacy of receiving vaccination alone, while Morimoto et al. [35] demonstrated the potential of infiltrating a Category III wound site with rabies vaccine as an alternative to the administration of RIG. Moreover, although a five-dose Essen regimen was considered for simplicity in the calculation of the direct costs of PEP, it has been indicated that the Japanese PCEC-K vaccine is less potent than those produced overseas [36]. The PEP regimen using the Japanese vaccine requires five to six intramuscular doses, i.e. a potential extra dose on Day 90, with clinical data suggesting that up to 85.4% of the patients (n=813) acquired satisfactory antibody titres after five vaccinations [20, 37]. Finally, anxiety associated with a dog bite that may develop into rabies has been suggested as an additional component in Years of Life lived with Disability (YLD) contributing to the economic burden of the disease, but it was not considered in the current model due to a lack of scientific validation of this assumption [15].

Results of scenario analysis demonstrated that the maintenance of current annual dog rabies vaccination policy could be economically efficient if several conditions were met. In worse-case analysis, the *BCR* would become above 1 if the risk of rabies introduction increased to 0.04 corresponding to a level of risk where rabies would enter Japan in 26 years while the economic burden of a rabies outbreak under the abolition of vaccination policy increased to $7.53 billion, a level close to the annual global burden of endemic canine rabies which was estimated to be $8.6 billion by Hampson et al. [15] (Table 3). The best-case analysis further illustrated that the economic efficiency of the current policy could be improved by decreasing the price of single rabies vaccination, relaxing the frequency of vaccination to every two to three years and implementing the policy on a smaller scale, e.g. only in targeted prefectures with the highest risk of rabies incursion (similar to the concept of immune belt, an example would be the one built by Malaysia along the border with Thailand) (Table 4). Overall, future research is highly warranted to provide further evidence-based information to determine whether the continuation of the current vaccination policy, as part of national rabies prevention system, is scientifically justified in the long run or not.

If the current policy were to be abolished, resources, specifically the recurrent funds provided by the government to JVMA for organization of annual vaccination campaign in all the 47 prefectures of Japan, could be allocated to more efficient uses to strengthen the national rabies prevention system. Based on the information from the investigated prefectural governments, a vaccination campaign in a particular prefecture with a capacity to vaccinate 12,000 to 14,000 dogs would receive financial support of about $173,665 and this suggests that with the abolition of the current policy a fund of around $12 to $14 could potentially be saved from each dog that would otherwise be vaccinated in the campaign. It should be noted that dog owners would still have the option to voluntarily vaccinate their dogs against rabies in private clinic even when vaccination campaign became unavailable under the abolition of the current policy. In terms of recommended reinforcement of the current rabies prevention system, simulation exercises of the contingency plan should be conducted regularly and continuous training of private veterinary clinicians and government officers in the rabies control team are very important [38]. Moreover, the current PEP delivery system must be strengthened in terms of the stockpile of human rabies vaccine and the emergency supply of RIG. Currently, there are approximately 114 local hospitals or clinics which offer rabies PrEP or PEP and about 40,000 to 50,000 Japanese human rabies vaccines are produced locally with a similar amount being imported each year [19, 2018 MHLW data]. Nevertheless, the local stockpile of human rabies vaccines appeared temporarily exhausted when there were two reports of introduced human rabies cases from the Philippines in November 2006 and the number of tourists seeking PEP after returning from overseas increased three-fold [19, 20]. Training of doctors and medical professionals is also essential to facilitate correct and efficient delivery of PEP to patients with real need. The potential use of PEP due to public panic would incur a substantial and unnecessary economic burden, emphasising that the importance of continuous public education to raise awareness and knowledge of rabies (Fig 2). Furthermore, the cost-effectiveness of PEP has been an international research topic and regimens consisting of fewer doses to reduce costs and fewer consultations to promote patient’s compliance have been examined [39, 40]. The use of Japanese PCEC-K vaccine in a three-dose intradermal PrEP regimen has been proved safe and efficacious [41, 42] (Shiota et al., 2008; Yanagisawa et al., 2012). Thus, further research on the suitability of the Japanese vaccine to time-and dose-sparing PEP regimens such as 4-dose Essen regimen, Zagreb regimen and one-week, 2-site ID regimen is highly warranted [39]. Lastly, it is expected that the active surveillance system on wild animals and the diagnostic capacity of reference laboratories in Japan will continue to be reinforced given the national guidelines of animal rabies survey was released in 2015.

## Conclusions

The implementation of the policy of mandatory annual vaccination of domestic dogs in rabies-free Japan is very economically inefficient for the purpose of reducing the economic burden of a potential canine rabies outbreak. Scenario analysis revealed that the economic efficiency of the current policy could be improved by decreasing the vaccination price charged to dog owners, relaxing the frequency of vaccination to every two to three years and implementing the policy on a smaller scale such as targeted prefectures instead of the whole Japan.

## Acknowledgements

We would like to thank Dr. Shoji Miyagawa from the Ministry of Health, Labour and Welfare, the prefectural government officials from Ibaraki, Tokushima and Miyazaki Prefectures and Dr. Satoshi Inoue from the National Institute of Infectious Disease for providing valuable information and comments to this study.

## Author Contributions

Conceptualization: Akio Yamada, Katsuagi Sugiura

Investigation and Methodology: Nigel C.L. Kwan, Katsuagi Sugiura

Data Curation, Formal Analysis, Software and Visualization: Nigel C.L. Kwan

Resources and Supervision: Katsuagi Sugiura

Validation: Akio Yamada, Katsuagi Sugiura

Writing – Original Draft Preparation: Nigel C.L. Kwan

Writing – Review & Editing: Nigel C.L. Kwan, Akio Yamada, Katsuagi Sugiura

